# A Simulation-Based Evaluation of Total-Evidence Dating Under the Fossilized Birth-Death Process

**DOI:** 10.1101/436303

**Authors:** Arong Luo, David A. Duchêne, Chi Zhang, Chao-Dong Zhu, Simon Y.W. Ho

**Author notes:** Correspondence: Arong Luo, Key Laboratory of Zoological Systematics and Evolution, Institute of Zoology, Chinese Academy of Sciences, Beijing 100101, China; Simon Ho, School of Life and Environmental Sciences, University of Sydney, Sydney, New South Wales 2006, Australia.

## Abstract

Bayesian molecular dating is widely used to study evolutionary timescales. This procedure usually involves phylogenetic analysis of nucleotide sequence data, with fossil-based calibrations applied as age constraints on internal nodes of the tree. An alternative approach is Bayesian total-evidence dating, which involves the joint analysis of molecular data from present-day taxa and morphological data from both extant and fossil taxa. Part of its appeal stems from the fossilized birth-death process, which provides a model of lineage diversification for the prior on the tree topology and node times. However, total-evidence dating faces a number of considerable challenges, especially those associated with fossil sampling and evolutionary models for morphological characters. We conducted a simulation study to evaluate the performance of total-evidence dating with the fossilized birth-death model. We simulated fossil occurrences and the evolution of nucleotide sequences and morphological characters under a wide range of conditions. Our analyses show that fossil occurrences have a greater influence than the degree of among-lineage rate variation or the number of morphological characters on estimates of node times and the tree topology. Total-evidence dating generally performs well in recovering the relationships among extant taxa, but has difficulties in correctly placing fossil taxa in the tree and identifying the number of sampled ancestors. The method yields accurate estimates of the origin time of the fossilized birth-death process and the ages of the root and crown group, although the precision of these estimates varies with the probability of fossil occurrence. The exclusion of morphological characters results in a slight overestimation of node times, whereas the exclusion of nucleotide sequences has a negative impact on inference of the tree topology. Overall, our results provide a detailed view of the performance of total-evidence dating, which will inform further development of the method and its application to key questions in evolutionary biology.

## Introduction

Resolving the evolutionary timescale of the Tree of Life has been one of the long-standing goals of biological research. There has been remarkable progress in this area over the past few decades, driven largely by analyses of genetic sequences using the molecular clock (Ho 2014; Donoghue and Yang 2016). Bayesian phylogenetic approaches hold particular appeal because they provide a unified framework for implementing models of nucleotide substitution, evolutionary rates, and lineage diversification (dos Reis et al. 2016; Bromham et al. 2018). At the same time, Bayesian molecular dating can incorporate calibrating information into the priors on node times. These calibration priors are usually applied to internal nodes of the tree, based on interpretations of relevant fossil evidence (Ho and Phillips 2009; Donoghue and Yang 2016) or biogeographic events (Ho et al. 2015b; De Baets et al. 2016). However, the recent introduction of total-evidence dating, in which fossils are included in the analysis as sampled taxa, enables the ages of the tips to supply the calibrating information (Pyron 2011; Ronquist et al. 2012).

Total-evidence dating makes use of both molecular sequence data and morphological characters, with fossil species being analysed together with their living relatives (e.g., Pyron 2011; Ronquist et al. 2012). In this approach, the phylogenetic positions of the fossil taxa are informed by the morphological characters and age information is provided directly by the fossils, without the need to specify calibration priors for internal nodes in the tree. This makes it possible to include all of the fossils available for the group being studied (Ronquist et al. 2012). In this regard, total-evidence dating differs from previous ‘node dating’ approaches, in which each calibration is typically based on only the oldest known fossil assigned to the clade descending from that node. A key advantage of total-evidence dating is that it obviates the need to specify any maximum age constraints, which are often chosen without strong justification but have potentially large impacts on the resulting date estimates (Hug and Roger 2007). A further benefit is that total-evidence dating eliminates the potential problem of marginal calibration priors differing from the user-specified priors, which arises when the latter are combined multiplicatively with the tree prior (Heled and Drummond 2012). Total-evidence dating has been used to infer the evolutionary timescales of various groups of taxa, including birds (Gavryushkina et al. 2017), fishes (e.g., Near et al. 2014; Arcila et al. 2015; Arcila and Tyler 2017), mammals (e.g., Slater 2013; Herrear and Davalos 2016; Kealy and Beck 2017), and plants (e.g., Larson-Johnson 2016).

An important step in the evolution of total-evidence dating was the development of the fossilized birth-death (FBD) process (Stadler 2010). This model is designed to generate the probability density of a tree with individuals sampled through time in an epidemiological or phylogenetic context. In phylogenetic analyses that involve total-evidence dating, the FBD process provides a model of lineage diversification that accounts for speciation, extinction, fossilization, and taxon sampling (Heath et al. 2014; Gavryushkina et al. 2014). The FBD model can allow sampled ancestors, whereby sampled fossils are direct ancestors of other taxa in the data set. Extensions of the FBD model include treating the process of fossilization and sampling as a piecewise function (Gavryushkina et al. 2014) and accommodating different taxon-sampling strategies (Zhang et al. 2016). More recent developments integrate multispecies coalescent models (Ogilvie et al. 2018), speciation modes (Stadler et al. 2018), or among-lineage variation in diversification rates (Mitchell et al. 2018). The FBD model can also be used without morphological characters, but under these circumstances the approach benefits from constraints on the placements of the fossil taxa (Heath et al. 2014). Alternatively, the FBD model can be used to analyse data sets that exclusively comprise morphological characters from fossil taxa and their extant relatives (Matzke and Wright 2016; Bapst et al. 2016; King et al. 2017; Matzke and Irmis 2018).

The FBD model has parameters that represent the speciation rate (*λ*), extinction rate (*μ*), and fossil recovery rate (*ψ*), along with the start time of the process (origin time *t_or_* or root age *t_mrca_*) and sampling fraction of extant taxa (*ρ*). For mathematical convenience, the model is parameterized using the net diversification rate (*d* = *λ − μ*), turnover rate (*r* = *μ/λ*), and fossil sampling proportion (*s* =*ψ*/(*μ+ψ*)) (Heath et al. 2014). Simulation-based studies have shown that the FBD model is generally able to recover the parameters used for simulation, though with some exceptions (Gavryushkina et al. 2014; Zhang et al. 2016); for example, the uncertainty in turnover rates has been found to vary with the size of the tree and with the sampling strategy. Drummond and Stadler (2016) found that the FBD model was able to infer the ages of fossil samples with a good degree of accuracy, confirming its internal consistency with other models within the Bayesian framework. However, analyses of empirical data have yielded date estimates that are often considerably younger when using the FBD model than when using other tree priors for total-evidence dating (Herrera and Dávalos 2016; Zhang et al. 2016; Gavryushkina et al. 2017). There have also been some discrepancies between the results of total-evidence dating and node dating (Vea and Grimaldi 2016; Arcila and Tyler 2017; Gustafson and Miller 2017; Kealy and Beck 2017).

In principle, total-evidence dating using the FBD model provides a satisfying approach because it combines the available data from both fossil and extant taxa. However, the inclusion of morphological data and fossil taxa presents a number of complex challenges. First, fossil specimens are often incomplete or fragmentary, leading to potential difficulties in resolving their phylogenetic placements (Sansom et al. 2010; Sansom and Wills 2013). Second, morphological data are typically analysed using the Mk model (Lewis 2001), a *k*-states generalization of the Jukes-Cantor model of nucleotide substitution (Jukes and Cantor 1969), but this model makes strong simplifying assumptions that are unlikely to hold true for real data. Third, morphological characters are likely to evolve in a far less clocklike manner than nucleotide sequences (dos Reis et al. 2016; Donoghue and Yang 2016; Drummond and Stadler 2016). Fourth, although the ages of fossils are usually treated as being known without error, this assumption is potentially problematic (O’Reilly et al. 2015). Finally, rates of fossilization and fossil sampling might vary across clades, whereas the FBD process typically assumes homogeneity of these rates throughout the tree (Matschiner et al. 2017).

In this study, we evaluate the performance of total-evidence dating under a range of conditions. Our analyses are based on synthetic data generated by simulating fossil occurrences, evolution of nucleotide sequences, and evolution of morphological characters on trees generated under a birth-death process. Using the FBD model for the tree prior, we examine how Bayesian estimates of node times and tree topologies are affected by fossil occurrences, number of morphological characters, and degree of among-lineage rate heterogeneity. In addition, we consider the influence of the model of morphological evolution and uncertainty in fossil ages. The results of our analyses allow us to present some practical guidelines for total-evidence dating using the FBD model.

## Materials and Methods

### Species Trees and Fossil Occurrences

Using TreeSim (Stadler 2011) in R (R Core Team 2017), we simulated speciation according to the birth-death process to produce 1000 trees (Stadler 2009), each with 50 extant species and between 8 and 83 extinct lineages (median = 31). These simulations were performed using a constant speciation rate *λ* = 0.05 per Myr, constant extinction rate *μ* = 0.02 per Myr, and sampling fraction *ρ* = 1 (i.e., complete sampling of present-day taxa). The diversification process was conditioned on the number of extant species, with speciation and extinction rates chosen to generate appropriate root ages and numbers of extinct tips. From the 1000 complete trees that were produced, we selected 20 trees that had crown ages of about 100 Ma (±1 Ma). These 20 trees varied in the total number of tips (74–105), origin times *t_or_* (101–196 Ma), and tree shapes as measured by the Colless index (3.7–8.5, corrected by the number of tips; Colless 1982). Ten of the 20 trees had root ages *t_mrca_* equal to their crown ages *t_c_*, whereas the others had root ages that were greater than their crown ages (Table 1; Figs. 1a and 1b).

**Figure 1.**
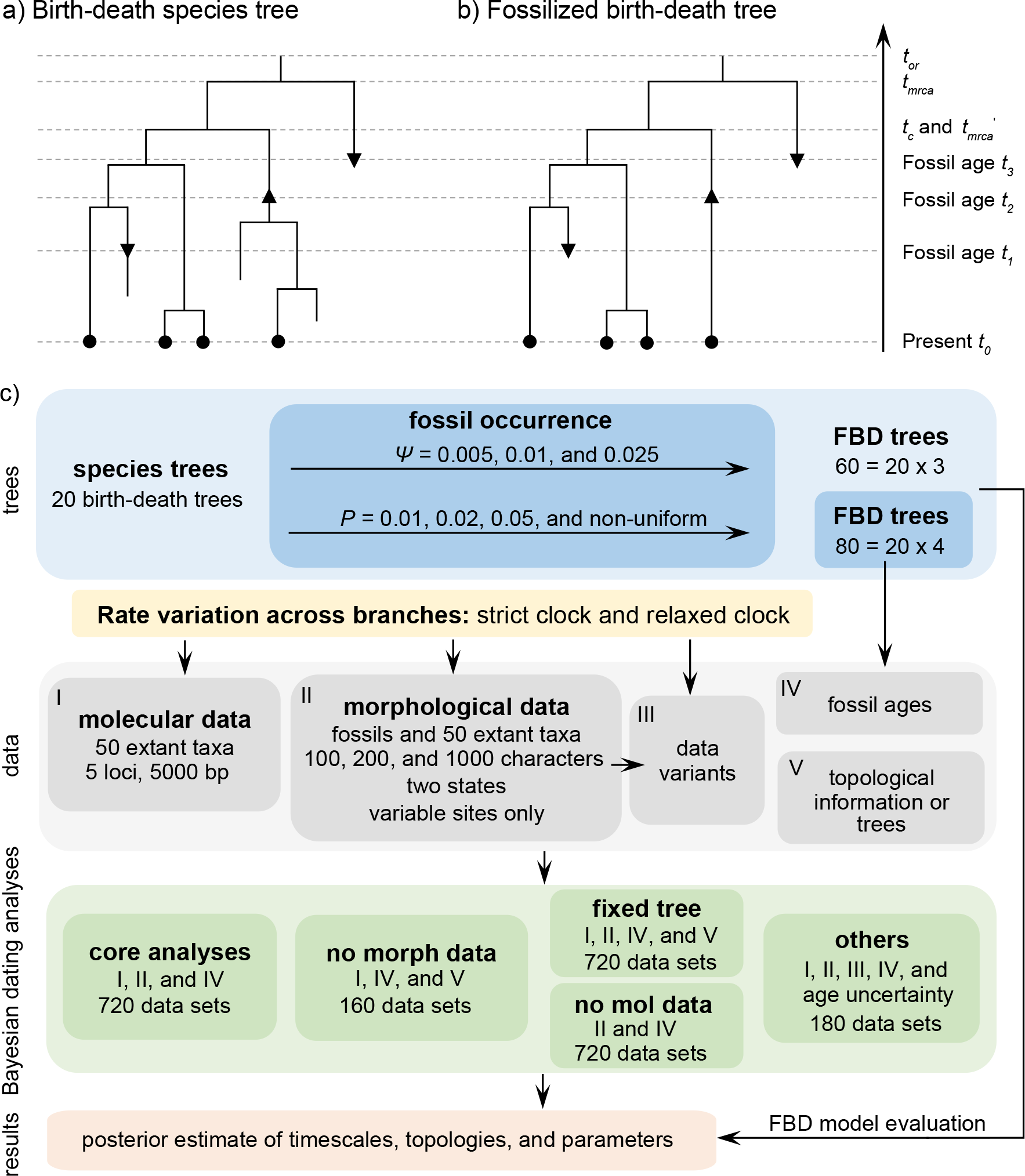
(a) Illustration of a complete tree generated under the birth-death process. Lineage diversification is controlled by birth rate *λ*, death rate *μ*, and sampling fraction *ρ*. From the origin time (*t_or_*) to the present day (*t_0_*), fossils have been sampled at *t_1_*, *t_2_*, and *t_3_*, with one (denoted by the upwards triangle) leaving an extant descendant (denoted by solid circle) and the other two (denoted by downwards triangles) leaving no extant descendants. With complete sampling of present-day taxa (*ρ* = 1), the age of the crown group *t_c_* remains the same, whereas the age of the root *t_mrca_* depends on whether fossils are sampled between *t_mrca_* and *t_mrca_*’. (b) The fossilized birth-death (FBD) tree depicting the reconstructed history of present-day taxa and sampled fossil taxa based on the complete species tree in (a). (c) Flow-chart showing the simulation pipelines and analyses conducted in this study. A detailed explanation of each step is provided in Materials and Methods. Briefly, we obtained the FBD trees by simulating speciation using the birth-death process, with the probability of fossil occurrences based on either *P* and *ψ*. Among these FBD trees, the 80 trees with fossil occurrences sampled by *P* were the main basis of this study. These trees provided the fossil ages and topologies. We simulated the evolution of nucleotide sequences and morphological characters on these trees, under various models of rate variation among lineages. We carried out series of Bayesian dating analyses under a range of settings and using various subsets of the data. These analyses yielded estimates of the posterior distribution of tree topologies, node times, and model parameters.

We used two different approaches to simulate fossil sampling on each of the 20 complete birth-death trees (Fig. 1c). First, we used a single parameter, *P*, to represent the probability of fossil occurrence; preservation potential and sampling intensity were not specified separately (Heath et al. 2014; Warnock et al. 2017). Fossil occurrences were modelled as a Bernoulli process in time slices of 2 Myr throughout the duration of each species tree, except along the branch between *t_mrca_* and *t_or_*. We employed three uniform models of fossil occurrence (*P* = 0.01, 0.02, and 0.05) across the 20 trees, then considered a simple non-uniform model in which *P* decreases linearly with *t* from 0.05 (*t* = 2) to 0.005 (*t* = 100) and ultimately to zero (Table 1).

In our second approach to simulating fossil sampling, we modelled the occurrence of fossils as a continuous process along each branch (except between *t_mrca_* and *t_or_*) of the 20 complete trees. We used three values for the fossil recovery rate (*ψ* = 0.005, 0.01, and 0.025), which characterizes a Poisson process of fossil recovery in the FBD model (Heath et al. 2014). These values were chosen to be half of those of *P* in the uniform models (because the time increments of *ψ* and *P* here are 1 Myr and 2 Myr, respectively), so that our two approaches to simulating fossil sampling were expected to generate similar numbers of fossil occurrences. Fossil occurrences sampled by this second approach were only used to evaluate the FBD process and to confirm the expected relationship between the values of *P* and *ψ*.

**Table 1.** Details of the 20 birth-death species trees and fossil occurrences sampled according to a parameter representing the probability of fossil occurrence *P*.

| Index<sup>1</sup> | Tree<br>1 | Tree<br>2 | Tree<br>3 | Tree<br>4 | Tree<br>5 | Tree<br>6 | Tree<br>7 | Tree<br>8 | Tree<br>9 | Tree<br>10 | Tree<br>11 | Tree<br>12 | Tree<br>13 | Tree<br>14 | Tree<br>15 | Tree<br>16 | Tree<br>17 | Tree<br>18 | Tree<br>19 | Tree<br>20 |
| --- | --- | --- | --- | --- | --- | --- | --- | --- | --- | --- | --- | --- | --- | --- | --- | --- | --- | --- | --- | --- |
| $t_{or}$ (Ma) | 101 | 102 | 103 | 105 | 105 | 105 | 106 | 107 | 110 | 113 | 122 | 132 | 141 | 142 | 148 | 159 | 161 | 166 | 175 | 196 |
| $t_{mrca}$ (Ma) | 99 | 99 | 100 | 100 | 101 | 104 | 100 | 101 | 100 | 100 | 100 | 131 | 121 | 140 | 119 | 100 | 154 | 161 | 173 | 185 |
| $t_c$ (Ma) | 99 | 99 | 100 | 100 | 99 | 101 | 100 | 101 | 100 | 100 | 100 | 99 | 100 | 100 | 101 | 100 | 100 | 100 | 100 | 99 |
| No. of tips (all) | 86 | 74 | 78 | 93 | 92 | 96 | 78 | 76 | 75 | 85 | 105 | 90 | 98 | 74 | 93 | 93 | 77 | 74 | 75 | 78 |
| Colless index (all) | 3.9 | 4.7 | 3.7 | 4.6 | 6.9 | 8.5 | 4.0 | 3.8 | 5.2 | 5.2 | 4.8 | 7.2 | 7.7 | 5.9 | 7.9 | 4.6 | 4.1 | 7.8 | 6.9 | 7.2 |
| Colless index (extant) | 2.1 | 2.0 | 2.7 | 3.2 | 2.3 | 3.9 | 2.9 | 2.4 | 3.9 | 4.9 | 3.6 | 2.8 | 2.5 | 2.9 | 3.0 | 2.6 | 2.5 | 2.8 | 2.8 | 2.8 |
| No. of fossils, $P = 0.01$ | 5 | 8 | 4 | 9 | 13 | 7 | 6 | 12 | 5 | 5 | 14 | 8 | 11 | 5 | 8 | 7 | 8 | 7 | 5 | 8 |
| No. of fossils, $P = 0.02$ | 13 | 12 | 16 | 18 | 24 | 17 | 21 | 16 | 17 | 17 | 29 | 14 | 24 | 12 | 10 | 19 | 21 | 15 | 20 | 13 |
| No. of fossils, $P = 0.05$ | 37 | 37 | 35 | 42 | 40 | 44 | 35 | 44 | 42 | 35 | 54 | 48 | 64 | 35 | 36 | 39 | 43 | 38 | 34 | 55 |
| No. of fossils, non-uniform $P$ | 30 | 18 | 28 | 41 | 37 | 20 | 33 | 29 | 31 | 20 | 50 | 30 | 37 | 24 | 31 | 38 | 26 | 24 | 25 | 30 |
| Max. age (Ma), $P = 0.01$ | 38 | 92 | 80 | 40 | 64 | 94 | 66 | 38 | 96 | 52 | 86 | 78 | 74 | 52 | 84 | 78 | 78 | 66 | 52 | 166 |
| Max. age (Ma), $P = 0.02$ | 60 | 66 | 94 | 72 | 96 | 102 | 80 | 42 | 66 | 78 | 96 | 88 | 118 | 136 | 114 | 68 | 142 | 102 | 144 | 178 |
| Max. age (Ma), $P = 0.05$ | 74 | 82 | 96 | 100 | 74 | 100 | 82 | 84 | 92 | 98 | 94 | 124 | 118 | 132 | 96 | 86 | 142 | 144 | 170 | 176 |
| Max. age (Ma), non-uniform $P$ | 92 | 64 | 78 | 64 | 62 | 84 | 60 | 74 | 58 | 74 | 78 | 62 | 72 | 78 | 82 | 66 | 78 | 56 | 96 | 76 |

<sup>1</sup>Note: $t_{or}$ , the origin time of a species tree; $t_{mrca}$ , the root age of a species tree; $t_c$ , the age of the crown group in a species tree; No. of tips (all), the total number of tips in a species tree; Colless index (all), the corrected Colless index by the number of all tips in a species tree; Colless index (extant), the corrected Colless index by the number of extant tips in a species tree; No. of fossils, the total number of fossils sampled in a species tree by the probability of fossil occurrence $P = 0.01, 0.02, 0.05$ , or non-uniform; Max. age, the maximum age of sampled fossils in a species tree by the probability of fossil occurrence $P = 0.01, 0.02, 0.05$ , or non-uniform.

### Simulations of Character Evolution

For each FBD tree, we simulated the evolution of nucleotide sequences along the reconstructed history of the extant species only. Two models of among-lineage rate variation were used to transform the chronograms into phylograms in NELSI v0.2 (Ho et al. 2015a). First, we assumed a strict molecular clock with a rate of 10^−3^ subs/site/Myr. Second, we used the white-noise model to allow rate variation across branches, with mean 10^−3^ subs/site/Myr and standard deviation 2×10^−4^ subs/site/Myr. Sequence evolution was simulated using Seq-Gen v1.3.4 (Rambaut and Grassly 1997) to produce five 1000 bp sequence alignments (equivalent to five ‘loci’) for each phylogram, with relative evolutionary rates randomly sampled from a symmetric Dirichlet distribution with α = 3. Simulations were performed using the HKY+G substitution model with base frequencies {A:0.35, C:0.15, G:0.25, T:0.25}, transition/transversion ratio *κ* = 4.0, and a gamma shape parameter of 0.5.

We simulated the evolution of morphological characters for both extant species and sampled fossils along each of the 20 FBD trees. To account for rates of morphological evolution being more likely to vary among lineages (dos Reis et al. 2016), we used a white-noise model of branch rates with a mean of 10^−3^ changes/character/Myr and three different standard deviations: 0 (i.e., a strict clock), 2×10^−4^ changes/character/Myr, and 5×10^−4^ changes/character/Myr. We used Seq-Gen first to simulate nucleotide sequence evolution with base frequencies {A:0.0, C:0.0, G:0.5, T:0.5}, then converted the resulting nucleotides into binary characters by recoding G to 0 and T to 1 (Puttick et al. 2017). Our simulations produced data sets of three sizes: *l* = 100, 200, and 1000 characters.

For our core analyses, which are described in detail below, nucleotide sequences from the extant species were combined with morphological characters from both the extant species and the sampled fossils. These produced three different scenarios of rate variation among lineages: nucleotide sequences under a strict clock with morphological characters under a strict clock (‘SS’); nucleotide sequences under a strict clock with morphological characters under moderate rate variation (‘SM’); and nucleotide sequences under moderate rate variation with morphological characters under high rate variation (‘MH’). Our simulations produced 240 data sets under each combination of clock models, differing with respect to their underlying FBD trees and/or the numbers of morphological characters.

### Total-Evidence Dating

#### Evaluation of the FBD Process

We first evaluated the outcomes of the FBD process with fossil occurrences sampled by *P* and *ψ* on the 20 birth-death trees, paving the way for the subsequent dating analyses. For each of the 140 FBD trees (Fig. 1c), we fixed the tree topology, branch lengths (in units of Myr), *t_or_*, and *ρ* to their true values, and excluded the sequence data and morphological characters in turn. We adopted diffuse priors, including beta distributions *B*(1,1) for the turnover *r* and fossil sampling proportion *s*, and an exponential distribution with mean 0.1 for the net diversification *d*. We estimated the posterior distribution using Markov chain Monte Carlo (MCMC) sampling with the SA (sampled ancestor) package in BEAST v2.4.8 (Bouckaert et al. 2014; Gavryushkina et al. 2014). Samples were drawn every 5000 steps over a total of 20 million steps and a burn-in fraction of 0.25. In the absence of the molecular and morphological data, these samples were effectively drawn from the prior distribution. During the MCMC sampling, we used a modified version of BEAST 2.4.8 that allowed us to retain the sampled ancestors, which were very sensitive to the automatic adjustments to branch lengths by the MCMC proposal mechanisms in BEAST. Sufficient sampling was checked using Tracer v1.7 (Rambaut et al. 2018).

#### Core Analyses

We used the FBD model for the tree prior in our analyses of the data sets produced by our simulations, with the ages of the sampled fossils treated as point values (Fig. 1c). The HKY+G model with four rate categories was used for the nucleotide sequences (Yang 1994), whereas the Mkv model was used for the variable characters in the morphological data (Lewis 2001). We applied separate uncorrelated lognormal relaxed-clock models to the molecular and morphological data (Drummond et al. 2006), with a uniform prior *U*(10^−6^,1) for the mean rate in all analyses. For the parameters of the FBD model, we used relatively diffuse priors: beta distributions *B*(1,1) for *r* and *s*; an exponential distribution with mean 0.1 for *d*; and a uniform distribution for *t_or_*, with a maximum bound of 300 Ma and a minimum bound matching the age of the oldest sampled fossil. The sampling proportion of extant species was fixed to 1 to match the settings used in our simulations.

The 720 data sets produced by our simulations were analysed using BEAST with the SA package. For each data set, we carried out two independent MCMC analyses in order to check for convergence. Each MCMC analysis consisted of 100 million steps, with samples drawn every 5000 steps and with a discarded burn-in fraction of 0.25. We checked for sufficient sampling by ensuring that all parameters had effective sample sizes of at least 100. Maximum-clade-credibility trees were identified from the combined samples using TreeAnnotator v1.8.4. To investigate the maximum-clade-credibility tree topology, we pruned the fossils to produce annotated trees containing only extant taxa. An additional MCMC analysis was performed without data, to allow us to evaluate the combined signal from the sequence data and morphological characters.

#### No Morphological Characters

The FBD model can be used for molecular dating in the absence of morphological characters, such that the diversification process is marginalized over all of the possible placements of the fossil occurrences (Heath et al. 2014). We performed analyses with the morphological characters excluded, so that the data comprised only the nucleotide sequences (generated using either a strict clock or with moderate among-lineage rate variation) and fossil occurrence times (i.e., 160 data sets in total). We used two different strategies for specifying the placements of the fossils in the tree during MCMC sampling. First, we imposed a monophyletic constraint for each fossil so that it was placed into its correct clade, as defined by its parent node. Second, we did not specify any constraints on the tree, so that we sampled full trees for the fossil positions conditioned on fixed *ρ* = 1 (Gavryushkina et al. 2014). Other settings for the MCMC analyses were the same as for the core analyses (Fig. 1c).

#### No Molecular Data

To examine the performance of Bayesian dating with the FBD model applied to morphological data sets, we excluded nucleotide sequences from the 720 data sets of the core analyses. Dating analyses were then performed using the morphological characters of the extant and fossil taxa, along with the fossil occurrence times. We used the same settings as for the core analyses, but with samples drawn every 1000 steps from a total of 20 million MCMC steps.

#### Fixed Tree Topology

We carried out total-evidence dating using the sequence data and morphological characters, with the FBD tree topologies fixed to those used for simulation. Clock models, choices of priors, and the MCMC settings were the same as those used for the core analyses. The sole exception was that posterior distributions of parameters were estimated from samples drawn every 2000 steps from a total of 40 million MCMC steps.

#### Alternative Conditions

We performed further analyses to investigate the influence of several factors that appeared to be influential in our core analyses. These analyses were all based on the sequence data and morphological characters produced by simulation with the SS pattern of among-lineage rate variation, fossil occurrences obtained using *P* = 0.05, and other relevant settings (as appropriate for each set of analyses). Given the three sizes of morphological character sets (i.e., *l* = 100, 200, and 1000) and 20 FBD trees, there were 60 data sets for each set of analyses (Fig. 1c).

First, we replaced the binary morphological characters with four-state morphological characters. To generate these data, we simulated the evolution of nucleotide sequences using Seq-Gen with the Jukes-Cantor model, then converted the nucleotides in the resulting sequences to numerical multistate coding (A to 0, C to 1, G to 2, and T to 3). Second, we performed dating analyses using the Mk model with the full sets of binary morphological characters, rather than using the Mkv model with only the variable characters. In our third set of analyses here, we aimed to test whether accounting for uncertainty in fossil ages would have any impact on date estimates (O’Reilly et al. 2015). We used uniform priors rather than point values for the fossil sampling times. The bounds of these uniform priors were chosen to match the boundaries of the stratigraphic stage from which each fossil was sampled, as defined by the International Commission on Stratigraphy (February 2017). For example, if a fossil had been sampled at 76 Ma, we instead used a uniform prior *U*(71.9,83.8) to reflect the age boundaries of the Campanian stage of the Upper Cretaceous. Other settings for the MCMC analyses were the same as those used for our core analyses.

### Evaluation of Total-Evidence Dating

Our main objectives were to examine the estimates of node times and/or tree topologies inferred from data generated under different simulation conditions. We thus treated estimates for the 20 birth-death species trees as independent replicates under each set of conditions (e.g., 20 repeats under the SS pattern of among-lineage rate variation, *P* = 0.05, and *l* = 100). Unless noted otherwise, we did not consider in any detail the differences across replicates.

To evaluate the performance of total-evidence dating with the FBD model, we examined the posterior medians of model parameters and node times. We chose not to focus on the posterior means, because this caused problems for identifying sampled ancestors in the maximum-clade-credibility trees. To allow comparisons of estimated node times, we used three metrics that involved standardizing the absolute estimates. First, we computed relative bias, which is the distance between the posterior median and the true value, divided by the true value. Second, we computed the relative 95% credibility interval (CI) width, which is the 95% CI width divided by the true (point) value. Third, we computed the coverage probability, which is the proportion of 95% CIs that contain the true values. Additionally, after pruning fossil taxa, we used the gamma statistic (Pybus and Harvey 2000) and stemminess rank (Fiala and Sokal 1985) to summarize relative node depths in the maximum-clade-credibility trees based on posterior medians of node times.

To evaluate the differences between each maximum-clade-credibility tree and the true topology, we used two measures of topological distance. First, we computed the absolute Robinson-Foulds topology distance, which is defined as twice the number of internal branches defining different bipartitions of the tips (Robinson and Foulds 1981; Penny and Hendy 1985). Second, we corrected this distance by the number of tips. Distance calculations were performed using the R package ape (Paradis et al. 2004; Popescu et al. 2012). To measure the performance of total-evidence dating in placing fossils into their correct phylogenetic positions, we split the sampled fossils into two categories, based on whether they had extant descendants or not (Figs. 1a and 1b). For fossils that left extant descendants, we measured whether their positions were correctly inferred or not by using two criteria: monophyletic grouping, which depends on whether the fossil is grouped with its extant and/or extinct relative(s); and being a sampled ancestor, which depends on whether its terminal branch length is zero or not. For fossils that did not leave extant descendants, we recorded whether they were correctly identified as sampled ancestors or not.

## Results

### Recovery of the FBD Parameters

The parameters of the FBD model were generally well recovered when the tree topology and branch lengths were fixed to those of the trees used for simulation (Fig. 2). As expected, our models with uniform *P* (*P* = 0.01, 0.02, and 0.05) led to fossil sampling proportions *s* similar to those from the three *ψ* values (*ψ* = 0.005, 0.01, and 0.025), confirming a mathematical approximation *P* ≈ 2 × *ψ* in our study. The median estimates of *s* across the 20 FBD trees approached the true values (0.2, 0.33, and 0.56, respectively). The model with non-uniform *P* yielded estimates of *s* most similar to those produced with *P* = 0.05, being generally consistent with the numbers of sampled fossils. The rates of net diversification (*d* = 0.03) and turnover (*r* = 0.4) were generally estimated accurately. However, we identified biases at finer scales, as seen in the estimated turnover rate increasing with *ψ* and *P*.

**Figure 2.**
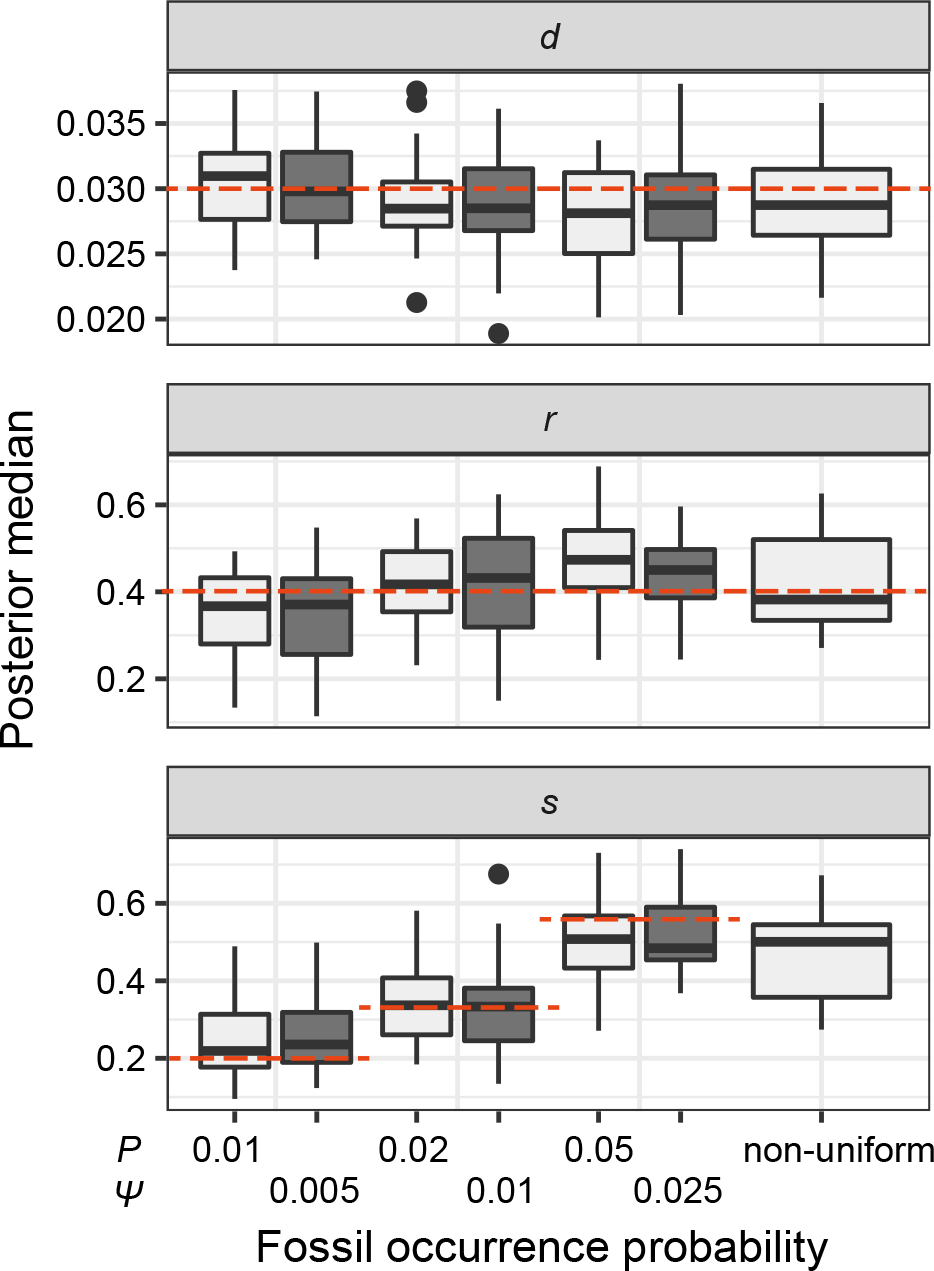
Posterior medians of the fossilized birth-death (FBD) model parameters from our evaluation of the FBD process while conditioning on fixed tree topologies and branch lengths. The three panels show boxplot summaries of posterior estimates of net diversification rate (*d* = *λ − μ*), turnover rate (*r* = *μ/λ*), and fossil sampling proportion (*s* = *ψ*/(*μ+ψ*)). Each summary is based on a set of 20 FBD trees, which were derived from fossil occurrences sampled by *P* (light grey shading) or *ψ*(dark grey shading) on our 20 simulated species trees. The dashed horizontal lines indicate the true values of *d*, *r*, and *s* that were used for simulation.

### Impacts of Rate Variation, Fossil Occurrences, and Number of Morphological Characters

Based on our core analyses, we examined the impacts of the three main factors that we varied across our simulations: degree of rate variation across branches, probability of fossil occurrence *P*, and number of binary morphological characters *l*. To evaluate their impacts via standardized metrics, we focused on the topological distance between the inferred topology and the true topology, as well as the date estimates for three key time points (Figs. 1a and 1b): the origin time of the FBD process (*t_or_*); the root age, or time to the most recent common ancestor of the sampled taxa (*t_mrca_*); and the crown age, or time to the most recent common ancestor of the extant taxa (*t_c_*).

**Figure 3.**
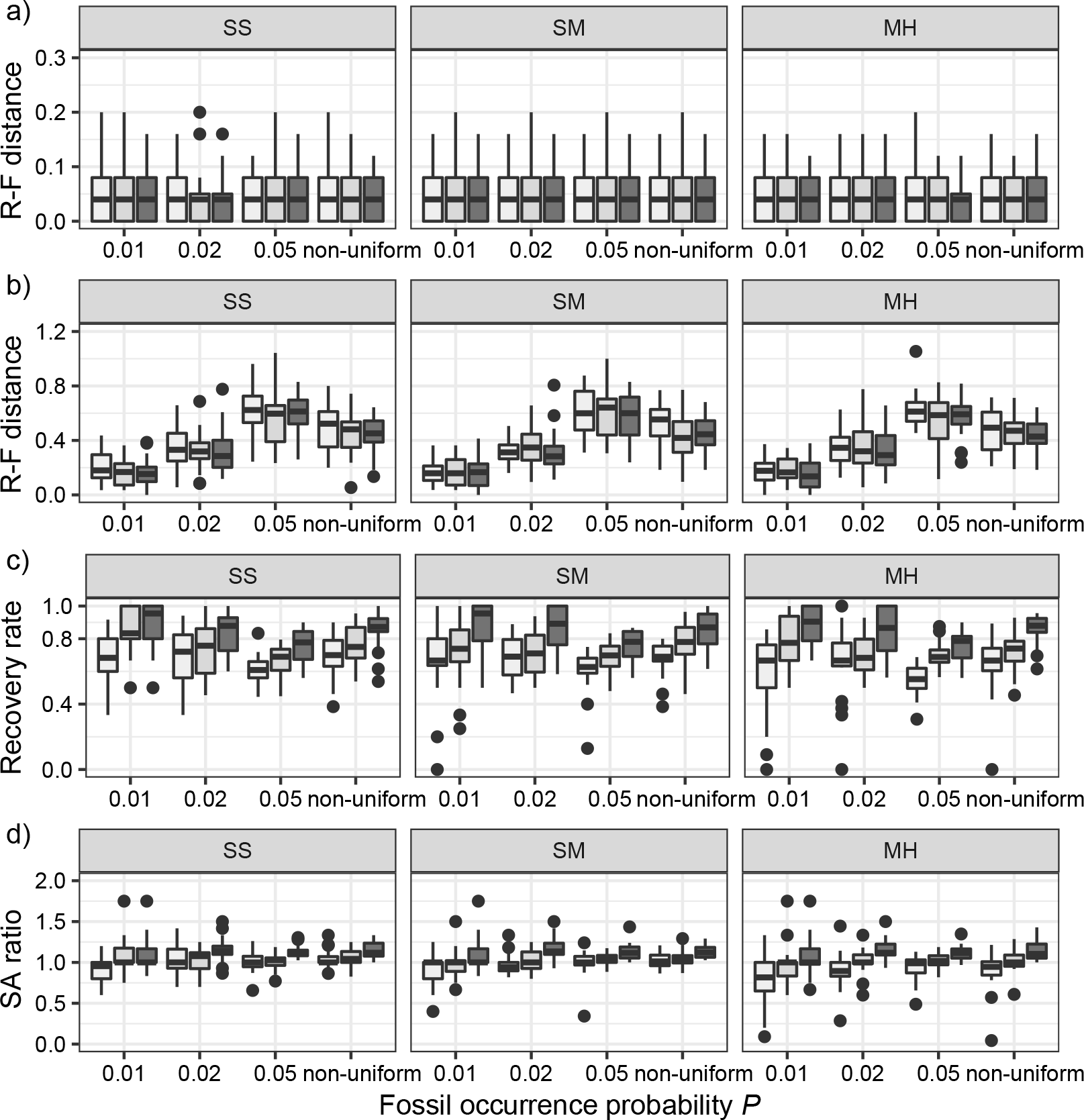
Performance of total-evidence dating in our core analyses in topological inference. Plots show corrected Robinson-Foulds (R-F) distances between maximum-clade-credibility trees and true trees while (a) excluding fossil taxa or (b) including fossil taxa. (c) Recovery rates of correct phylogenetic positions for fossil taxa that have left extant descendants. (d) Ratios of placing all sampled fossils as sampled ancestors (SA) in maximum-clade-credibility trees to the true numbers of sampled ancestors. Each panel shows the results from a different model of among-lineage rate variation for the molecular and morphological data: strict clock and strict clock (SS); strict clock and moderate rate variation (SM); and moderate rate variation and high rate variation (MH). Within each panel, boxplot summaries are shown for the 20 FBD trees under each model of fossil occurrence probability (*P* = 0.01, 0.02, 0.05, and non-uniform). For each fossil occurrence probability, results are shown for three different sizes of morphological characters (*l* = 100, 200, 1000 from left to right, in increasingly dark shades of grey).

The performance of topological inference depended on whether or not the sampled fossils were taken into account. When fossil taxa were pruned, the maximum-clade-credibility trees were very similar to those used for simulation (median of corrected Robinson-Foulds distances = 0.04; Fig. 3a). When fossil taxa were retained, however, the differences between the maximum-clade-credibility trees and true trees were much larger (median of corrected Robinson-Foulds distances > 0.1 if *P* = 0.01); corrected Robinson-Foulds distances increased with *P* in the models with uniform probabilities of fossil occurrence, while those for the analyses of data generated with the non-uniform *P* model fell between the values found in analyses with *P* = 0.02 and *P* = 0.05 (Fig. 3b). Topological distances showed weaker trends with the degree of rate variation and *l*, especially when the fossil taxa were pruned (Figs. 3a and 3b).

Coverage probabilities for *t_or_*, *t_mrca_*, and *t_c_* were 86.9%, 85.7%, and 82.1%, respectively. They did not show clear associations with the three main factors across the results of our core analyses, except for a lower coverage probability for *t_or_* when the probability of fossil occurrence *P* was non-uniform. We thus focus on relative bias and relative 95% CI width as respective measures of the accuracy and precision of our time estimates. We found that low rate variation across branches (SS or SM patterns of rate variation) led to estimates that had slightly better accuracy and precision than those from scenarios with higher rate variation across branches (MH pattern of rate variation; Fig. 4). The impacts of the fossil occurrence probability *P* and the number of morphological characters *l* varied among simulation treatments.

The accuracy of date estimates was not substantially affected by *P* or *l* (Fig. 4a), with the relative biases of *t_or_*, *t_mrca_*, and *t_c_* being close to 0. However, estimation accuracy for *t_or_* slightly increased with *P* in the uniform models. For the non-uniform *P* model, the spread of date estimates across the simulation replicates was similar to that when *P* = 0.01. There were some large positive biases in estimates for the three time points, although the proportions of estimates that were greater than the true dates were 58.6%, 58.8%, and 57.2% for *t_or_*, *t_mrca_*, and *t_c_*, respectively. Combinations of *P* and *l* had clear impacts on the precision of time estimates (Fig. 4b). For *t_or_*, *t_mrca_*, and *t_c_*, the relative 95% CI widths decreased with increasing *P* in the uniform models, while those with the non-uniform *P* model were smaller than those when *P* = 0.02 but larger than those when *P* = 0.05. As expected, the precision of time estimates generally increased with *l*. The relative 95% CI widths of the estimates of *t_or_* were generally greater and more variable across replicates than those of *t_mrca_* and *t_c_*. For example, given *P* = 0.01, means of the relative 95% CI widths were 0.78, 0.35, and 0.34 for *t_or_*, *t_mrca_*, and *t_c_*, respectively.

**Figure 4.**
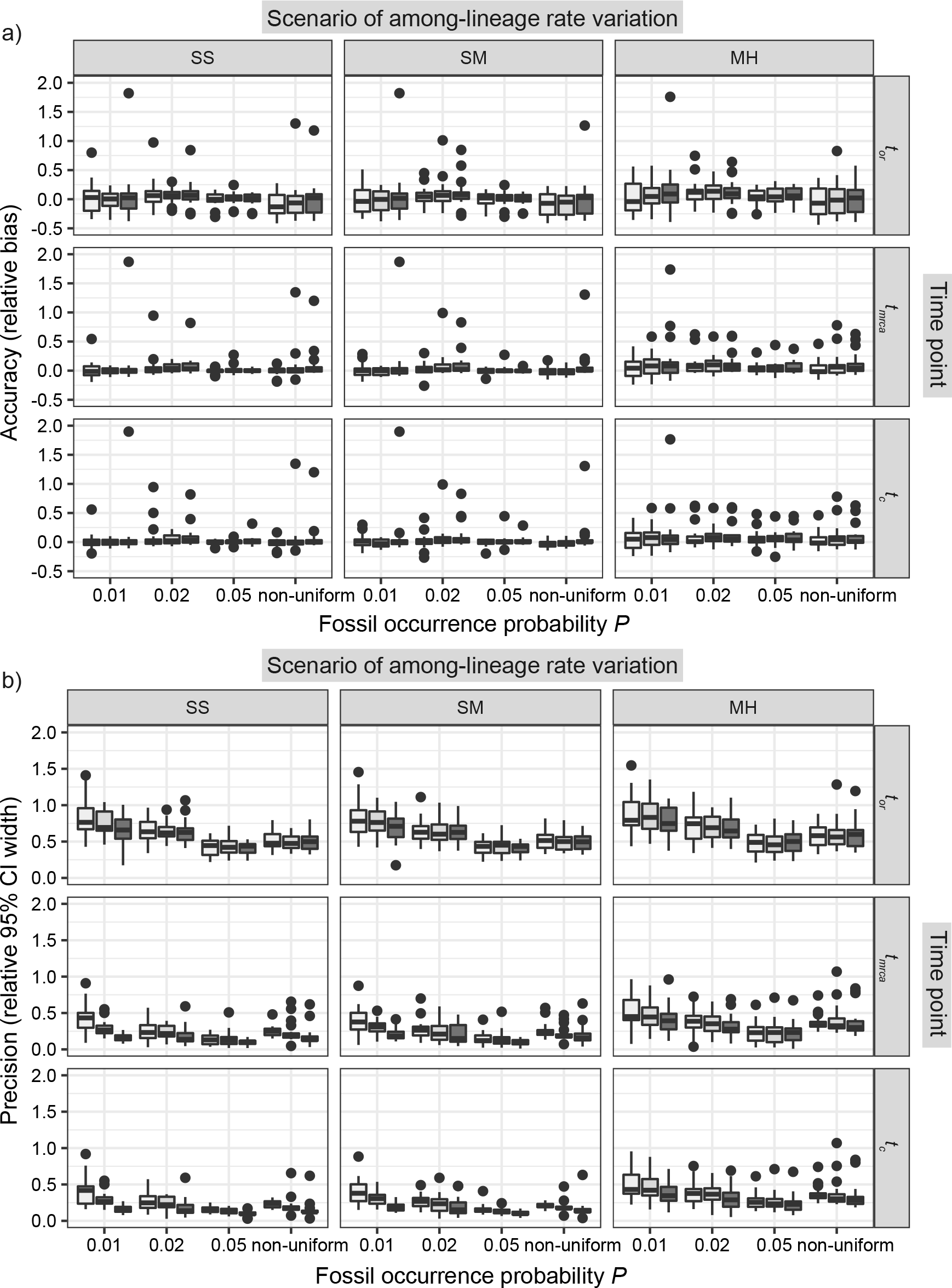
Performance of total-evidence dating in our core analyses in estimating origin time (*t_or_*), root age (*t_mrca_*), and crown age (*t_c_*). (a) Accuracy of estimates, as measured by relative bias (distance between posterior median and true value, divided by the true value). (b) Precision in estimates, as measured by relative 95% credibility interval (CI) width (posterior 95% CI width divided by the true value). Each column of panels shows the results from a different model of among-lineage rate variation for the molecular and morphological data: strict clock and strict clock (SS); strict clock and moderate rate variation (SM); and moderate rate variation and high rate variation (MH). Within each panel, boxplot summaries are shown for the 20 FBD trees under each model of fossil occurrence probability (*P* = 0.01, 0.02, 0.05, and non-uniform). For each fossil occurrence probability, results are shown for three different sizes of morphological characters (*l* = 100, 200, 1000 from left to right, in increasingly dark shades of grey).

### Relative Node Times and Placements of Fossil Taxa

We used the gamma statistic and stemminess rank to summarize the relative node times in the maximum-clade-credibility trees without fossils. When these were plotted against the corresponding metrics for the trees used for simulation, the lines of best fit had slopes close to 1.00 for all scenarios of rate variation. However, some biases were apparent with higher levels of among-lineage rate heterogeneity, as seen in the MH pattern of rate variation (Pearson’s correlation coefficient *R* = 0.93). This result was consistent with the outcomes of the date estimation described above. To examine the date estimates in further detail, we inspected the estimates for the youngest and median nodes. These nodes were chosen from the trees used for simulation and we ensured that they were present in the maximum-clade-credibility trees. Posterior medians of the ages of the two nodes were close to the true values whether the fossils were pruned or not. However, the date estimates for the youngest node in the tree had smaller biases than those for the nodes with median ages.

The large topological distances when the fossil taxa were retained in the maximum-clade-credibility trees revealed the difficulty in placing fossils correctly. Fossils with extant descendants were usually placed in the expected phylogenetic positions. Their recovery rates increased with *l*, but decreased with increasing probability of fossil occurrence in the models with uniform *P*. For the non-uniform *P* model, the recovery rates were similar to those when *P* = 0.02 (Fig. 3c). Among all sampled fossils, the ratios of placing them as sampled ancestors to the true numbers of sampled ancestors were greater than 1.0 for 368 out of 720 cases, a bias that increased with *l* (Fig. 3d). Thus, for the fossils without extant descendants, the numbers of sampled ancestors were generally overestimated, while absolute numbers of sampled ancestors being placed incorrectly tended to increase with *l*. These numbers of sampled ancestors were obtained from the maximum-clade-credibility trees, but the numbers increased considerably when based on the posterior medians from MCMC samples (the rate of being sampled ancestors >1.0 for 63.5% of cases).

We further evaluated potential differences across the species/FBD trees, with reference to the age estimates for key nodes (i.e., *t_or_*, *t_mrca_*, and *t_c_*). With respect to *l* and different scenarios of among-lineage rate variation, we treated estimates under these conditions as independent replicates. Thus, we had nine repeats for each of the 80 FBD trees. We found that the large positive biases in the date estimates tended to occur for three particular species trees (Trees 6, 10, and 11; Table 1), which were among the four trees that had the most imbalanced topologies for extant taxa (corrected Colless index > 3.5). However, this pattern was not always clear, especially when *P* = 0.05, and was only moderate for the relative 95% CI widths. The accuracy and precision of the estimates of *t_mrca_* and *t_c_* were broadly similar across the other species trees, for each value of *P*. For *t_or_*, however, relative biases and relative 95% CI widths were found to decrease with true *t_or_*, with the magnitude of these changes varying across the *P* models.

When fossils were pruned, distributions of topological distances were uneven across species trees. The absolute Robinson-Foulds distances ranged from 0 (e.g., Trees 8 and 9) to 10 (Trees 3 and 14). We did not carry out further comparisons when the fossils were retained, because they were sampled randomly on each species tree.

### Effects of Excluding Morphological Characters

We performed two sets of analyses without morphological characters, either with or without constraints on the placements of the fossil taxa. When we used monophyly constraints to restrict the placements of the fossil taxa, inference of the tree topology showed similar performance to the core analyses. However, there were slight improvements in the placement of fossil taxa, as reflected by smaller topological distances between inferred and true topologies (e.g., when *P* = 0.05, median of corrected Robinson-Foulds distances was < 0.3 compared with > 0.6 for the core analyses). The overestimation of the number of sampled ancestors was somewhat mitigated, with 96 out of 160 cases yielding sampled-ancestor rates not exceeding 1.0. Most of the fossils that left extant descendants were correctly placed (mean recovery rate 79.7% overall).

The accuracy of the posterior medians for *t_or_*, *t_mrca_*, and *t_c_* was poorer than when morphological data were included, although with fewer instances of extreme positive biases (when relative bias > 1.0). There was a greater tendency for the posterior medians to exceed the true values, which occurred in 65.0%, 75.0%, and 83.8% of analyses for *t_or_*, *t_mrca_*, and *t_c_*, respectively (Fig. 5a). Accuracy was slightly poorer when the sequence data had evolved with a moderate degree of rate variation among lineages. Coverage probabilities remained quite high overall, however, with the 95% CIs containing the true values between 78% and 91% of instances for the three time points. Compared with the analyses that included morphological data, there was also a reduction in the precision of the date estimates (Fig. 5b). Using the gamma statistic and stemminess rank to summarize the overall estimates of relative node times, the inferred values generally matched those for the trees used for simulation. Posterior medians for the ages of the youngest and median nodes in the maximum-clade-credibility trees were close to the true values, but the lines of best fit had slightly greater slopes than in the core analyses.

**Figure 5.**
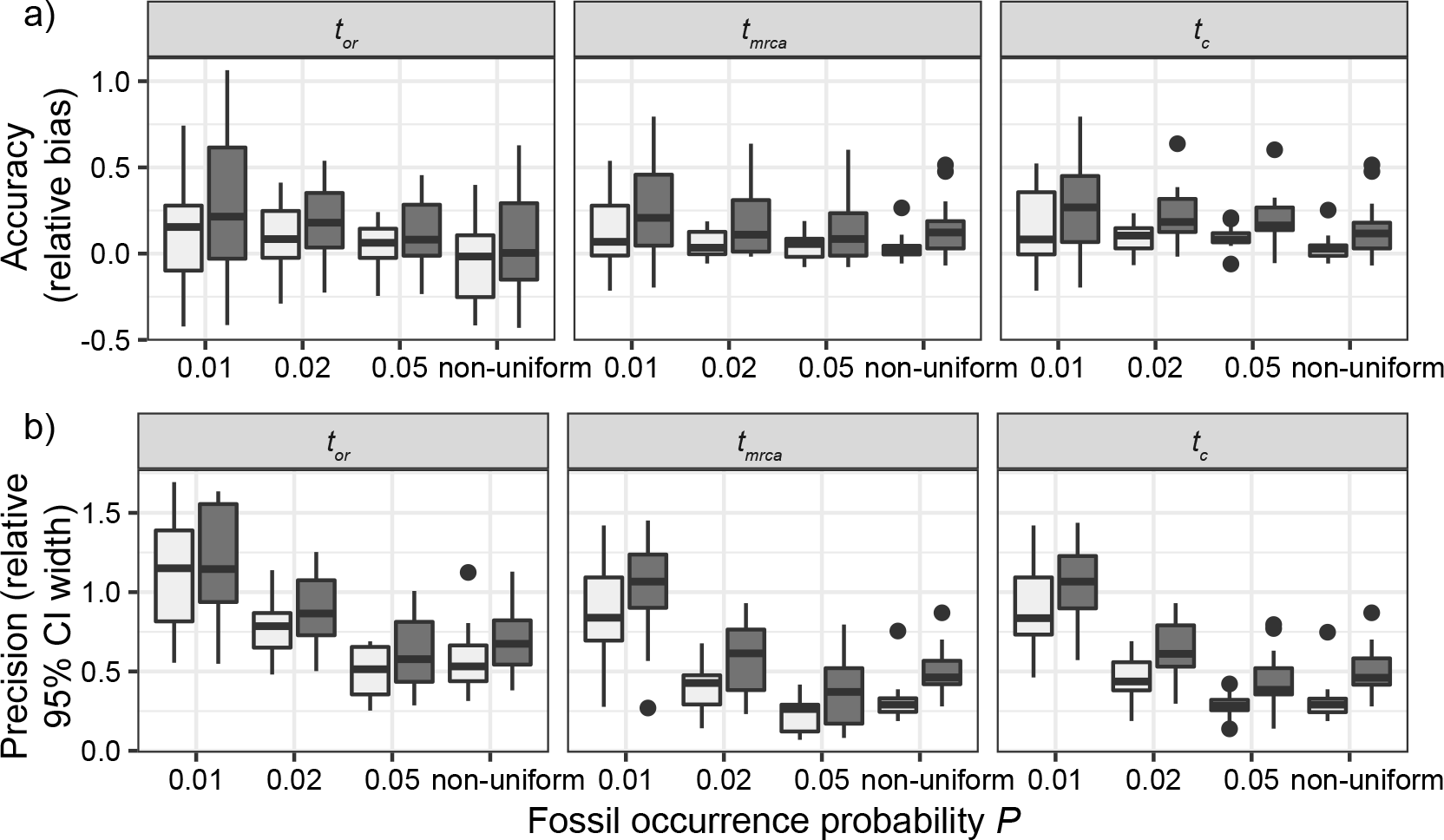
Posterior estimates for origin time (*t_or_*), root age (*t_mrca_*), and crown age (*t_c_*) in the analyses when morphological data were excluded. For each fossil occurrence probability (*P* = 0.01, 0.02, 0.05, and non-uniform), the left boxplot (light grey shading) shows estimates for molecular data that have evolved under a strict clock, whereas the right boxplot (dark grey shading) shows estimates that have evolved under moderate rate variation across branches. (a) Accuracy of estimates, as measured by relative bias. (b) Precision in estimates, as measured by relative 95% credibility interval width.

For the analyses in which morphological data were excluded and in which we did not specify any constraints on the tree topology, we experienced substantial problems with MCMC mixing and found that independent MCMC replicates did not converge. These problems were most pronounced when the data included large numbers of fossil taxa. Consequently, we do not report the detailed results of these analyses here.

### Effects of Excluding Molecular Data

We carried out analyses of the morphological data, with the nucleotide sequence data excluded. Topological inferences were generally similar to those of the core analyses. When fossils were pruned from the maximum-clade-credibility trees, corrected Robinson-Foulds distances from true topologies did not vary clearly across the *P* models (Fig. 6a). When fossils were retained, topological distances were larger and increased with *P* (Fig. 6b). In contrast with the results of the core analyses, however, larger *l* greatly reduced the topological distances whether the fossils were retained or not. Nevertheless, the distance estimates were greater overall than in the core analyses. For example, the overall median of the corrected Robinson-Foulds distances was 0.40 when fossils were pruned, ten times larger than that for the core analyses. Summaries of fossil positions in the maximum-clade-credibility trees did not differ much from those of the core analyses.

The accuracy and precision of the posterior estimates for *t_or_*, *t_mrca_*, and *t_c_* were generally consistent with those when nucleotide sequences were included, with coverage probabilities of 90.6%, 89.6%, and 83.8% for *t_or_*, *t_mrca_*, and *t_c_*, respectively. However, the summarized relative node depths via the gamma statistic and stemminess rank had greater biases under all degrees of rate variation, though the lines of best fit still had slopes close to 1.00. Whether the fossils were retained or not, posterior medians of the age estimates for median nodes were close to the true values. In contrast, there was a tendency to overestimate the ages of the youngest nodes in the trees (lines of best fit with slopes around 2.0).

**Figure 6.**
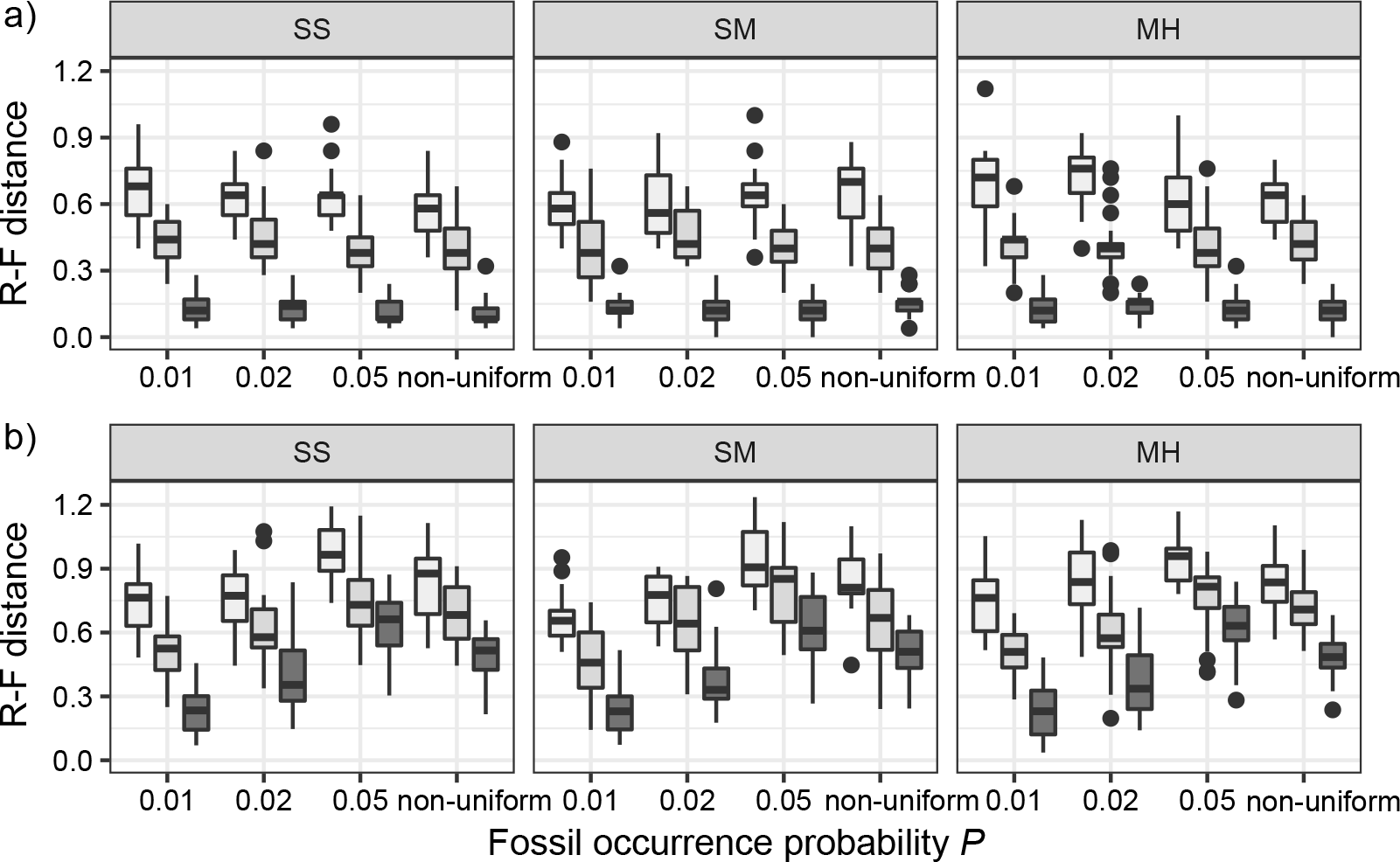
Posterior estimates in topological inference when molecular data were excluded. (a) Corrected Robinson-Foulds distances between maximum-clade-credibility trees and true trees with fossil taxa excluded. (b) Corrected Robinson-Foulds distances with fossil taxa included. Each panel shows the results from a different model of among-lineage rate variation for the molecular and morphological data: strict clock and strict clock (SS); strict clock and moderate rate variation (SM); and moderate rate variation and high rate variation (MH). Within each panel, boxplot summaries are shown for the 20 FBD trees for each model of fossil occurrence probability (*P* = 0.01, 0.02, 0.05, and non-uniform). For each fossil occurrence probability, results are shown for three different sizes of morphological characters (*l* = 100, 200, 1000 from left to right, in increasingly dark shades of grey).

### Effect of Fixing Tree Topology

We carried out analyses with both the morphological and molecular data, while fixing tree topologies during MCMC sampling. The overall ratio of sampled ancestors in maximum-clade-credibility trees to the true numbers of sampled ancestors was close to 1.00. In light of the fact that not all fossils that left extant descendants were recovered as sampled ancestors, overestimation of the number of sampled ancestors was still problematic sometimes for the fossils that did not have extant descendants. Estimation accuracy and precision for *t_or_*, *t_mrca_*, and *t_c_* were similar to those from the core analyses, except in two respects: there were almost no estimates with extreme relative biases; and the relative 95% CI widths were slightly smaller for *t_mrca_* and *t_c_*. Estimates for the summarized relative node depths, youngest nodes, and median nodes were generally close to the true values.

### Variations on the Conditions of the Core Analyses

**Figure 7.**
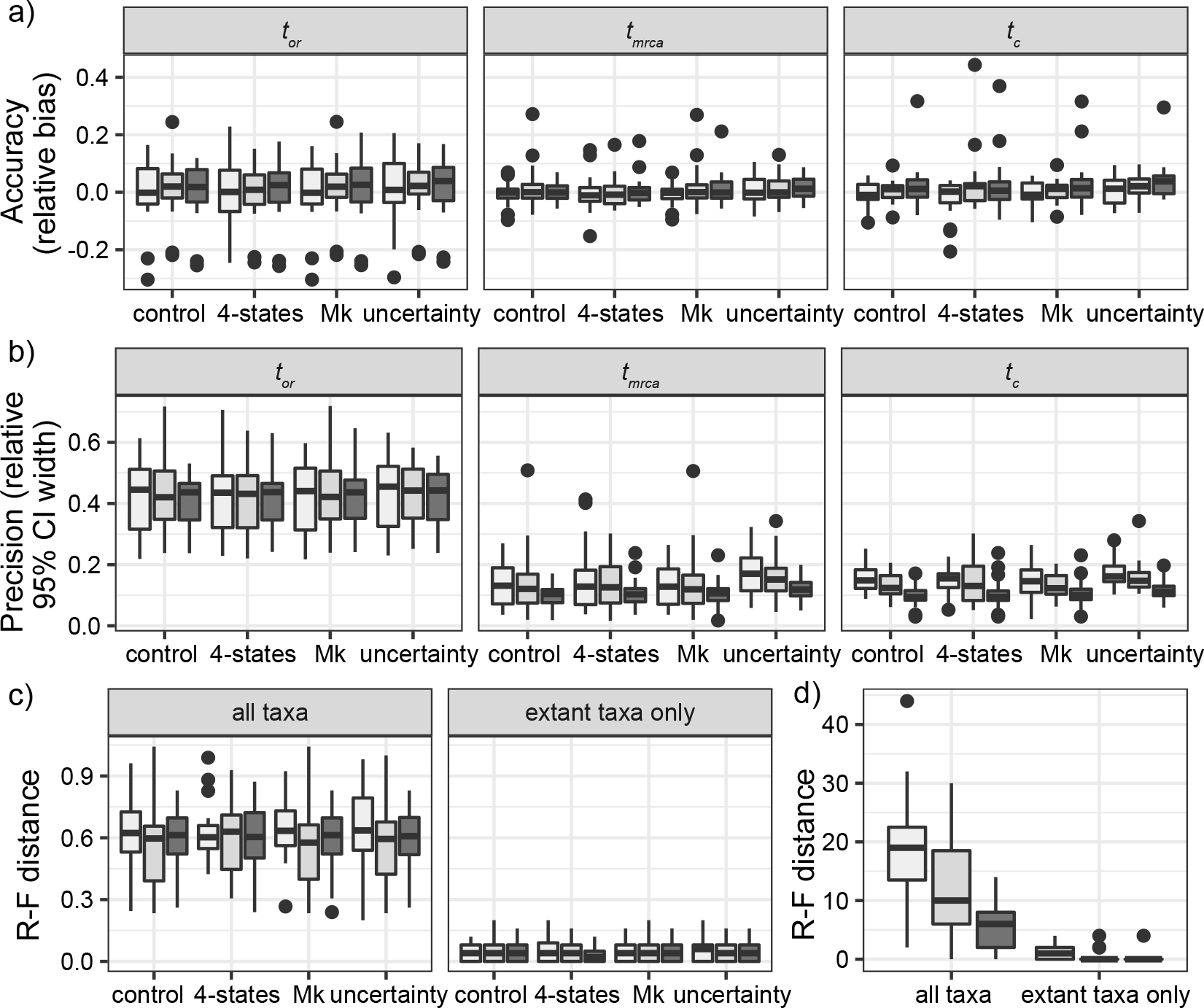
Performance of total-evidence dating under variations on the conditions of the core analyses. Results are shown for: counterpart analyses in the core analyses (denoted by ‘control’); those when binary morphological characters were replaced by 4-state morphological characters (denoted by ‘4-states’); those when the Mk model was used to analyse the full morphological data sets, rather than using the Mkv model to analyse only the variable morphological characters (denoted by ‘Mk’); and those when the uncertainty in fossil ages was taken into account (denoted by ‘uncertainty’). (a) Accuracy of posterior medians, as measured by relative bias, for origin time (*t_or_*), root age (*t_mrca_*), and crown age (*t_c_*). (b) Precision in date estimates, as measured by relative 95% credibility interval width. (c) Corrected Robinson-Foulds distances between the maximum-clade-credibility trees and the trees used for simulation. (d) Absolute Robinson-Foulds distance between maximum-clade-credibility trees derived from control analyses and those derived from the analyses taking into account fossil age uncertainty, based on either all taxa or only extant taxa. For each of the four treatments within each panel in (a), (b), and (c) and for the two treatments in (d), boxplots summarize the results for three different sizes of morphological characters (*l* = 100, 200, 1000 from left to right, in increasingly dark shades of grey).

We examined three variations on the conditions of the core analyses, with their counterparts from the core analyses being treated as the controls. First, we replaced the binary morphological characters with 4-state morphological characters. Second, we used the Mk model to analyse the full sets of morphological characters rather than using the Mkv model to analyse only the variable morphological characters. Neither of these variations led to any appreciable impacts on the accuracy and precision of date estimates, nor on the estimates of the tree topology (Fig. 7). In contrast, our third variation on the conditions of the core analyses, which involved incorporating uncertainty in the fossil sampling times, resulted in slightly wider relative 95% CI widths in estimates for *t_mrca_* (Fig. 7b). In this scenario, the inferred topologies of extant taxa did not differ substantially from the results of the core analyses. However, numerous discrepancies appeared when fossils were retained in the trees, and the absolute Robinson-Foulds distances decreased with *l* (Fig. 7d).

### Estimates of FBD Model Parameters

Estimates of the net diversification rate (*d*), turnover rate (*r*), and fossil occurrence probability (*s*) varied across the different conditions for simulation and analysis explored in this study (Fig. 8). When the tree topologies were fixed, posterior medians were similar to those from the aforementioned model recovery in our evaluation of the FBD process. In other cases, *r* tended to be underestimated and *s* tended to be overestimated, whereas the patterns for *d* were less consistent.

**Figure 8.**
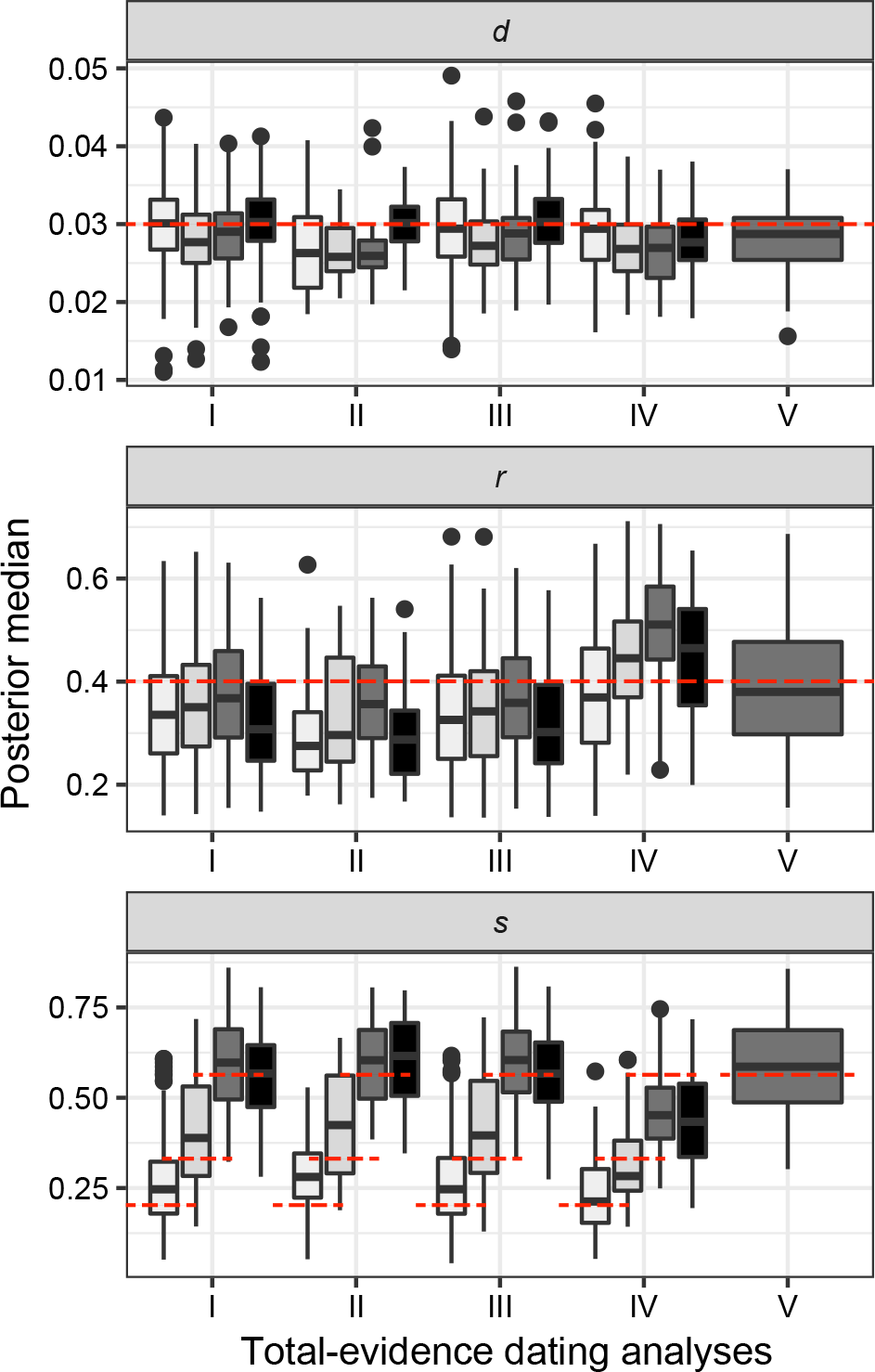
Posterior medians of the fossilized birth-death model parameters net diversification rate (*d*), turnover rate (*r*), and fossil sampling proportion (*s*) from all dating analyses. Results are shown for the core analyses (I), analyses without morphological characters (II), analyses without nucleotide sequences (III), analyses conditioned on fixed tree topologies (IV), and analyses under other variations on the conditions of the core analyses (V). Boxplots summarize the estimates from analyses grouped according to the *P* models used for simulation (*P* = 0.01, 0.02, 0.05, and non-uniform from left to right, in increasingly dark shades of grey). The red dashed lines indicate the true values of *d*, *r*, and *s* that were used for simulation.

The degree of evolutionary rate variation across branches did not produce clear impacts on the estimates of the three FBD parameters. However, the number of morphological characters (*l*) had a small impact in analyses conditioned on fixed tree topologies, but much larger effects in the other related analyses where *d* and *s* increased and *r* decreased with *l*. Compared with the results of our core analyses, using 4-state morphological characters or using the Mk model to analyse the morphological data had no apparent impact on the estimates of the FBD parameters. When we incorporated uncertainty in the fossil ages, however, *s* and *r* were slightly overestimated and underestimated, respectively.

## Discussion

### Joint Estimation of Tree Topology and Node Times

Our simulation study provides a range of insights into the performance of Bayesian total-evidence dating with the FBD model. We found that divergence times were accurately estimated under most of the conditions investigated in our core analyses, with relative biases being close to zero and posterior medians approaching the true ages. Relative 95% CI widths were usually below 1.0, indicating a moderate degree of precision in the divergence-time estimates. Our use of these measures of performance is consistent with those in previous studies (e.g., Gavryushkina et al. 2014; Zhang et al. 2016), but different from the use of other metrics such as coverage probability (Heath et al. 2014) and absolute 95% CI widths (Warnock et al. 2017).

The results of our core analyses revealed that the estimates for the origin time of the FBD process (*t_or_*) were less accurate and less precise than those for the ages of the root (*t_mrca_*) and the crown group (*t_c_*). This is consistent with expectations, given that taxa were not sampled during the interval between *t_mrca_* to *t_or_*, and that we used a diffuse prior for *t_or_*. The association between estimates of *t_or_* and other factors, such as the probability of fossil occurrence *P* and number of morphological characters *l*, was also different from those seen for *t_mrca_* and *t_c_*. These results highlight the difficulty in estimating *t_or_*, even under the most benign conditions explored in this study. However, this parameter is rarely of direct interest in total-evidence dating analyses, where there tends to be a much greater focus on the age of the root or the crown group.

The ages of deep nodes were overestimated in a moderate proportion of cases in our core analyses. This suggests that total-evidence dating with the FBD model is not particularly susceptible to ‘deep-root attraction’, a problem associated with unreasonably ancient divergence-time estimates (O’Reilly et al. 2015; Ronquist et al. 2016). In this respect, our results are consistent with the findings of previous studies (Herrera and Dávalos 2016; Zhang et al. 2016; Gavryushkina et al. 2017). However, we did observe some cases in which there were large positive biases in the age estimates for deep nodes. With reference to the results from differences in date estimates across the species/FBD trees and analyses with fixed tree topologies, these cases appeared to be the combined outcome of incorrect fossil placement and tree imbalance. This result partly echoes the findings of previous investigations of the impact of tree shape (Duchêne et al. 2015). Unfortunately, tree imbalance and problematic fossil placements are difficult to avoid in practice. Using informative priors that place a penalty on unobserved ghost lineages might help to mitigate the impacts of deep-root attraction on total-evidence dating (see Ronquist et al. 2016).

In Bayesian phylogenetic dating, the inferred evolutionary relationships are often also of interest. We found that total-evidence dating performed well in terms of inferring the relationships among extant taxa, which underscores the role of molecular data in providing a strong phylogenetic signal for these taxa. In contrast, the most notable failure of total-evidence dating is the incorrect phylogenetic placement of fossil taxa in the maximum-clade-credibility trees, as observed in our analyses. This outcome illustrates the challenges of topological inference and the resolution of deep nodes based on morphological characters, as identified in previous work (Puttick et al. 2017). We found that errors were reduced by increasing the number of morphological characters, but not by switching from binary to multistate character coding or including invariable sites.

Our simulations of the evolution of morphological characters involved a number of simplifications (Goloboff et al. 2018; O’Reilly et al. 2018). For example, we assumed independence among characters, a simple model of character replacement, and relatively simple patterns of among-lineage rate variation (Wright et al. 2016). In reality, however, morphological traits are subject to a range of selective pressures and potentially display differing modes and degrees of evolutionary rate heterogeneity across lineages (Lee and Palci 2015). Moreover, morphological characters are often very incompletely coded for fossil taxa (Sansom et al. 2010; Sansom and Wills 2013), but we did not include missing data in our simulations. Despite these simplifications, the key assumptions involved in our simulations were matched by those in the methods used to analyse the morphological data. The design of our study allowed us to avoid potential problems arising from model misspecification and model inadequacy, which are likely to be important problems for analyses of empirical data (Ronquist et al. 2016).

### Impacts of the Probability of Fossil Occurrence

A key benefit of total-evidence dating is that it allows the fossil record to be used more effectively in the estimation of evolutionary rates and timescales. However, the quality and completeness of the fossil record is subject to variations in depositional environments, taphonomy, time depth, and sampling intensity (Donoghue and Benton 2007; Holland 2016). Our simulations greatly simplified the process of fossil occurrence by ignoring the distinction between preservation potential and sampling intensity. Instead, the heterogeneous and incomplete nature of the fossil record was reflected in our use of different models of the probability of fossil occurrence, based on a single parameter *P*. The impact of this parameter on the estimates of divergence times and the tree topology was found to outweigh the effects of the other factors explored in our simulation study, such as the degree of among-lineage rate variation.

Our different models of the probability of fossil occurrence imparted multiple aspects of fossil occurrences along lineages in a birth-death species tree, which were associated with some distinct patterns among our results. With an increasing number of sampled fossils, the estimates of divergence times improved in precision. In our models with a low or non-uniform probability of fossil occurrence, there was a smaller chance of sampling older lineages and older fossils. This effectively led to a reduction in the range of sampled fossil ages. In turn, there was a negative impact on the accuracy of estimates of the origin time *t_or_* for the FBD process, but not of the age estimates for the root and crown group. Additionally, varying *P* can change the distribution of fossil occurrences on the tree topology, because imbalanced topologies tend to a greater number of long terminal branches. The limited number of replicates and the stochasticity of the fossil sampling procedures in our study preclude further exploration and interpretation of this effect.

The probability of fossil occurrences had no apparent impacts on the inference of the relationships among extant species. This outcome is presumably due to the rich information content of the nucleotide sequences in our simulations. A somewhat unexpected pattern, however, was that phylogenetic inferences tended to become worse overall with increasing numbers of fossils, due to the difficulty in placing fossil taxa correctly in the maximum-clade-credibility trees. For fossils without extant descendants, in particular, we found a tendency to overestimate the number of sampled ancestors. This potentially increases the number of ghost lineages across the full species tree. These fossils sometimes clustered in a separate group in the maximum-clade-credibility tree, with their phylogenetic placements partly reflecting their relative ages (results not shown; O’Reilly et al. 2015; Donoghue and Yang 2016).

When we accounted for uncertainty in the ages of the fossil occurrences, we found a measurable decline in the precision of the divergence-time estimates. This differs from the results of previous analyses based on tip-dating methods, which found that incorporating uncertainty in the ages of ancient DNA samples did not noticeably affect the estimates of the root age (Molak et al. 2013). Moreover, we found that uncertainty in fossil ages affected the placement of these fossils in the maximum-clade-credibility trees, which supports the notion that the placement of fossils should be informed by both their ages and their morphological characters (O’Reilly et al, 2015; Lee and Yates 2018). Our investigation of the observed impacts of fossil-age uncertainty is based on the relatively narrow age ranges of the stratigraphic stages to specify the uncertainty in fossil sampling times. In reality, however, age uncertainty can be much greater because of uncertainties in radiometric dating, biocorrelation, and other factors.

### Dating with Restricted Data Sets or Conditions

The full potential of total-evidence dating with the FBD prior is realized in joint analyses of morphological and molecular data sets, but the method can also be used when only one of these types of data is available. In the absence of morphological data, the fossil occurrence times can be used to inform the FBD model (Heath et al. 2014). Although we found that this approach has the potential to mitigate the problem of deep-root attraction (Ronquist et al. 2016), the inclusion of monophyly constraints on the fossil occurrences was essential to the tractability of the dating analyses. Even with optimal constraints on monophyly, however, the divergence times tended to be overestimated. This was possibly because the fossil occurrences were still able to jump between placements on the stem lineage or into the crown group defined by the monophyly constraints. When a stem fossil is incorrectly placed in the crown group, the age of the crown group will tend to be overestimated. The extent of age overestimation was exacerbated by the presence of rate variation across branches.

The FBD model is increasingly being used to analyse data sets comprising only morphological characters (e.g., Bapst et al. 2016; Matzke and Irmis 2018). We found that the exclusion of molecular data led to large reductions in the performance of phylogenetic inference, which confirms the substantial challenges facing Bayesian phylogenetic inference with morphological characters alone. Our method of tree summarization using the maximum-clade-credibility topology might have led to apparently worse performance than using majority-rule-consensus topologies (O’Reilly and Donoghue 2018), but we chose to use the former for the sake of consistency across our analyses. Our results pointed to some decoupling of the inferences of divergence times and tree topology, given that date estimates for key nodes remained accurate even while the quality of topological inference declined. However, we did observe greater biases in estimates of relative node times, along with overestimation of the age of the youngest node in the tree.

When we fixed the tree topology, we found a substantial improvement in the performance of total-evidence dating. There were far fewer cases of extremely biased estimates of node times, while the number of sampled ancestors for all sampled fossils was correctly recovered. The greatest improvements in performance were seen for highly imbalanced trees. However, given that the true tree topology is almost never known in practice, our results do not necessarily provide support for the sequential inference of the tree topology and divergence times when compared with a joint estimation procedure (in contrast with O’Reilly et al. 2015).

### Macroevolution and the Fossilized Birth-Death Process

The FBD process provides a convenient tree prior for Bayesian analyses of combined paleontological and neontological data, but can itself also provide valuable information about the diversification rates of the lineages being studied. Our results from the evaluation of the FBD process showed good accuracy in recovering the parameters of the FBD model: the net diversification rate *d*, turnover rate *r*, and fossil sampling proportion *s*. However, perhaps partly due to the use of diffuse priors for these parameters, we found some unexpected variation in estimates of *d* and *r* for different probabilities of fossil occurrence. Biases in estimation of the FBD model parameters are potentially problematic because they can mislead interpretations of the macroevolutionary process. For example, using the formula *λ* = *d*/(1 *− r*) and *μ* = *rd*/(1 *− r*) (Heath et al. 2014), we found both speciation and extinction rates were somewhat underestimated across our analyses (overall means were 0.04 and 0.01, respectively). Among all of the dating analyses that we carried out, only those with fixed tree topologies yielded estimates of the FBD model parameters that were comparable to those obtained from our initial evaluations of the model. These results draw attention to the influence of the phylogenetic placements of fossil taxa and the numbers of sampled ancestors on the estimates of the FBD model parameters.

## Conclusions

Total-evidence dating offers a powerful means of understanding macroevolutionary processes, especially when coupled with the fossilized birth-death process as a tree prior in Bayesian analysis. Using a simulation-based approach, we have performed a comprehensive evaluation of the performance of Bayesian total-evidence dating with the FBD process. Our results have demonstrated that the evolutionary relationships of extant taxa are well estimated, while the precision of divergence-time estimates tended to increase with the number of sampled fossils. However, we encountered considerable difficulty in identifying the correct phylogenetic placements of fossil taxa, even with good sampling of morphological characters.

Our study has revealed the considerable challenges posed by the absence of morphological data when analysing a combination of extant and fossil taxa. Even though the FBD model can be used to infer evolutionary timescales using fossil occurrence times alone (Heath et al. 2014), the date estimates were sensitive to the presence of rate variation across branches even when topological constraints were applied to all fossils.

Overall, the results of our simulation study have demonstrated the general utility of the FBD model in total-evidence dating. Further studies involving comprehensive analyses of empirical data sets will provide deeper insights into the performance of these methods when using morphological and molecular data that have evolved under more complex conditions. Continued development and extension of the FBD model will help to unlock the potential of using the combined information in morphological, molecular, neontological, and paleontological data for resolving evolutionary timescales across the diversity of life.

## Acknowledgements

We acknowledge the University of Sydney’s high-performance computing cluster, Artemis, for providing computing resources that have contributed to the research results reported in this paper.

